# antigen-prime: Simulating coupled genetic and antigenic evolution of influenza virus

**DOI:** 10.64898/2026.01.23.701420

**Authors:** Zorian T. Thornton, Thien Tran, Marlin D. Figgins, John Huddleston, Trevor Bedford, Frederick A. Matsen, Hugh K. Haddox

## Abstract

Seasonal influenza virus undergoes rapid antigenic drift to escape population immunity. Computational methods can be used to organize viral genetic diversity into antigenically similar variants and estimate variant-specific growth rates. However, benchmarking these methods is challenging because it can be difficult to accurately quantify antigenicity and growth rates in nature. Simulating viral evolution using defined selective pressures can provide ground-truth data for benchmarking. But, existing simulators do not link genetic sequences to antigenic phenotypes under selection from host populations.

Here, we present a forward-time epidemic simulator called `antigen-prime` that links these factors. We use it to simulate viral evolution over 30 years and validate the simulation recapitulates genetic and antigenic patterns observed in natural influenza evolution.

We then use the simulated data to benchmark methods for assigning variants and estimating their growth rates. We evaluated a sequencebased and a phylogenetics-based method for variant assignment, finding the former was slightly more effective at separating viruses into antigenically distinct groups. We also evaluated methods for estimating variant growth rates in one-year sliding windows. Estimates were accurate in most windows, but highly inaccurate in several others. Examining high-error windows revealed several examples of a previously unreported failure mode. In all, `antigen-prime` provides a simulation framework to benchmark models of influenza evolution, and could be used to help guide future development of these models. The source code is openly available at https://github.com/matsengrp/antigen-prime.

## 1 Introduction

Influenza virus infects about one billion and kills hundreds of thousands of people globally each year (Krammer et al., 2018). Vaccination provides the primary method of prevention, but the virus undergoes rapid antigenic evolution, necessitating annual vaccine updates (Petrova and Russell, 2018). To track the virus’s evolution, laboratories worldwide sequence tens of thousands of influenza genomes each year, sharing data through public databases (Hadfield et al., 2018; Shu and McCauley, 2017). Public health agencies also monitor influenza spread through case counts and hospitalizations.

It is of interest to develop computational methods to use the above sources of data to help guide vaccine updates (Morris et al., 2018). One goal of such methods is to partition observed viral sequences into groups (i.e. “variants”) with similar antigenic phenotypes. Another goal is to estimate variant-specific growth rates, helping to quantify variant fitness in the present and forecast future variant abundance.

For seasonal influenza virus, two main computational methods exist for variant assignment. One method involves building a phylogenetic tree of observed sequences and then partitioning the tree into clades that correspond to unique variants (Neher et al., 2025). This method can incorporate prior knowledge about which mutations are most likely to impact the virus’s antigenicity. Another method involves computing pairwise distances between observed sequences, and then using dimensionality reduction to embed sequences in a low-dimensional space. Clustering of sequences in this space can then be used to identify unique variants (Nanduri et al., 2024). Both methods have proven useful for interpreting real-time influenza surveillance data (Huddleston et al., 2024; Nanduri et al., 2024).

A variety of methods exist for estimating variant-specific growth advantages from viral surveillance data. Some only consider variant-specific frequencies over time, such as the fitness model by Luksza and Lässig (Luksza and Lässig, 2014) or multinomial logistic regression approaches (Ito et al., 2021; Obermeyer et al., 2022; Piantham et al., 2021). Others also consider observed numbers of case counts over time, such as the fixed growth advantage (FGA) and growth advantage random walk (GARW) models from the evofr framework (Figgins and Bedford, 2025).

Although these methods are widely used, it is challenging to benchmark their accuracy. That is because natural populations lack known ground-truth variant assignments and growth advantages. Experiments such as neutralization assays or hemagglutination inhibition assays can be used to quantify antigenic phenotypes of influenza viruses in a laboratory setting. However, it can be difficult for experiments to fully capture antigenic selection in nature due to the high level of heterogeneity in human immune responses to influenza (Kikawa et al., 2025).

A complementary approach is to simulate viral evolution under defined selective pressures and then use the simulated data, and associated ground-truth quantities, to benchmark the above methods. Doing so would require a simulator that models three key aspects of influenza evolution: (1) genetic sequence evolution through a biologically realistic mutation processes, (2) coupled antigenic evolution where mutations in epitope regions alter a virus’s antigenic phenotype, allowing escape from host immunity, and (3) sustained viral transmission in a large host population, with case counts tracked over time. While numerous pathogen-evolution simulators exist (Bedford et al., 2012; Jariani et al., 2019; Moshiri et al., 2019; Ochsner et al., 2025), none fully integrate all three of these components. SANTA-SIM (Jariani et al., 2019) models genetic sequence evolution, but not antigenic phenotypes or transmission dynamics. Conversely, antigen (Bedford et al., 2012) models transmission dynamics and antigenic evolution, but does not model genetic sequence evolution. FAVITES (Moshiri et al., 2019) models contact network transmission without antigenic evolution, while virolution (Ochsner et al., 2025) models within-host evolution without population-level antigenic drift.

Here, we develop a simulator called `antigen-prime` that models all three components listed above. We use it to simulate 30 years of seasonal influenza evolution, verifying that the simulated data reproduce influenza-like dynamics across genealogical, antigenic, and epidemiological dimensions. We then use the simulated data to benchmark methods for assigning variants and predicting variant growth rates. While the methods perform well overall, we identify several examples where they perform poorly, exposing previously unrecognized failure modes that occur even with abundant data.

## 2 Results

### 2.1 Summary of antigen-prime

We sought to build a simulator that integrates all three components of influenza evolution listed above. We achieved this by extending the antigen simulator (Bedford et al., 2012), which already models transmission dynamics and antigenic evolution. We updated antigen to include explicit genetic sequences that evolve through mutation and drive antigenic change. We call this updated simulator `antigen-prime`.

The original antigen implements a deme-structured SIR model to simulate viral transmission dynamics and antigenic evolution under selective pressure from host immunity. Each infected host carries a single virus object that is associated with coordinates that represent the virus’s location in a twodimensional antigenic space. The Euclidean distance between two viruses in this space approximates their antigenic similarity, that is, the degree of immune cross-reactivity between them, with smaller distances corresponding to greater cross-reactivity. Mutations to the virus move it in this space. Individual hosts maintain immune memory as a collection of antigenic coordinates from previous infections. When an infected host comes in contact with a susceptible host, the minimum Euclidean distance in antigenic space between the virus from the infected host and viruses from the susceptible host’s immune memory determines the infection risk, with smaller distances conferring greater protection. Selection to escape immune memories in the host population drives viral antigenic evolution. However, viruses in antigen are not associated with genetic sequences as they exist only as points in antigenic space.

In `antigen-prime`, we updated the virus object to include not only the virus’s coordinates in antigenic space, but also a nucleotide sequence of a protein-coding gene (Figure 1A). Simulations initialize with a single virus with a defined sequence, and this sequence evolves as the virus replicates and spreads in the host population. The user must define a subset of amino-acid sites in the protein to be “epitope sites”, where mutations have large antigenic effects (all remaining sites are considered to be “non-epitope sites”). In this paper, we used an influenza hemagglutinin (HA) nucleotide sequence that codes for a protein of length 566 amino acids. We classified 49 of these amino-acid sites to be epitope sites, using the set of epitope sites defined in Luksza and Lässig (Luksza and Lässig, 2014).

**Figure 1:**
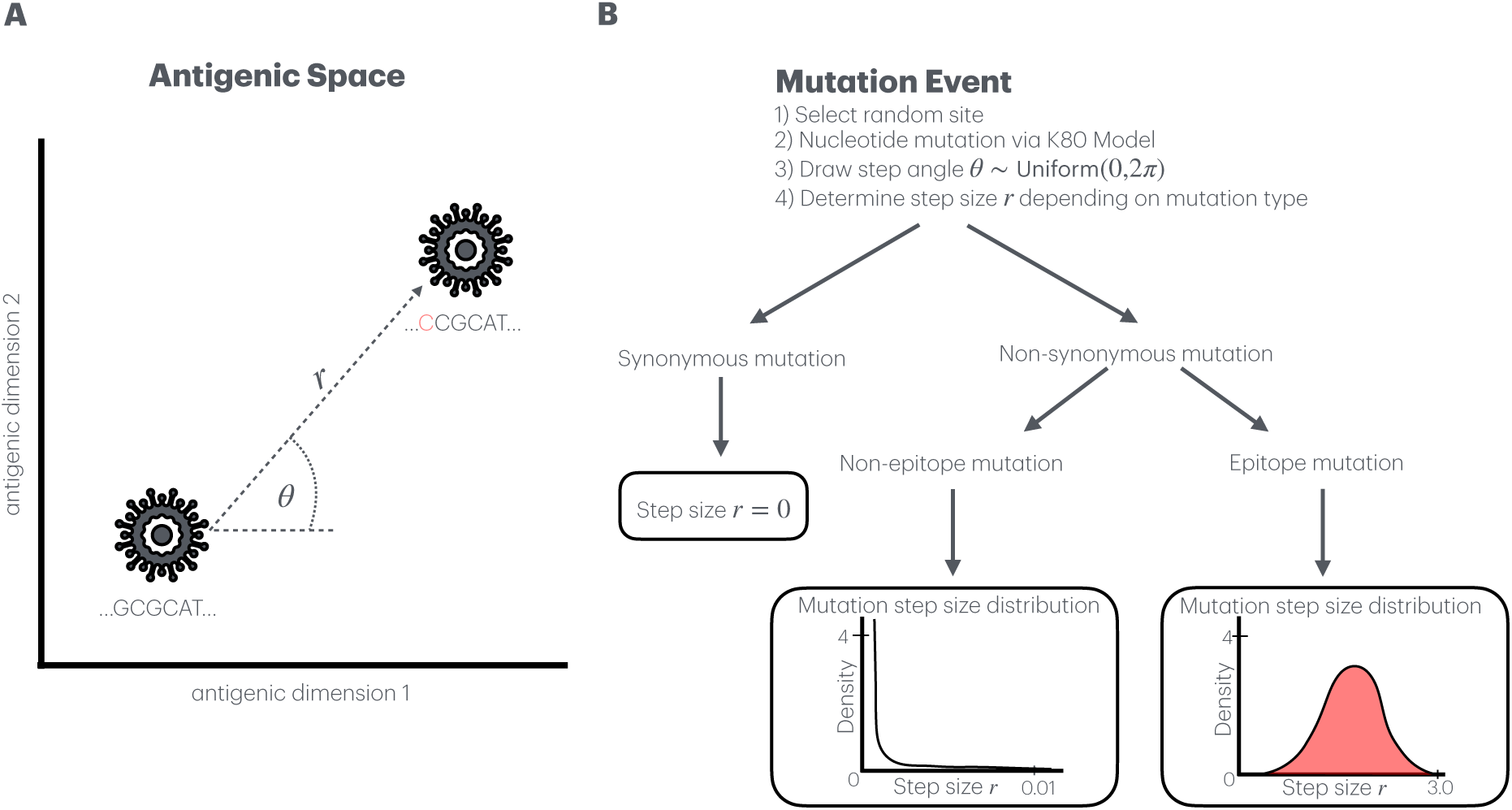
Schematic of how `antigen-prime` simulates coupled genetic and antigenic evolution of viruses. **A**: Each virus is represented by both a nucleotide sequence and coordinates in a two-dimensional antigenic space. Mutation events simultaneously change the nucleotide sequence and move the virus in antigenic space. **B**: During mutation events, a nucleotide site is randomly selected and mutated according to the K80 model, while antigenic coordinates are updated based on a sampled step direction *θ* and step size *r* that depends on whether the mutation is synonymous or non-synonymous and whether it occurs in an epitope or non-epitope site.

The nucleotide sequence of a virus object mutates at a user-defined mutation rate. Since there is only one virus object per infected host, this object’s sequence is like a consensus sequence within the host, and the rate at which it mutates represents the rate at which mutations reach sufficient frequency within the host to transmit to new hosts upon contact events. Thus, we model this rate as a function of both neutral mutation and within-host purifying selection. We model the neutral mutation process using the K80 mutation model (Kimura, 1980), parameterized by a transition/transversion ratio *κ* = 5.0 (user-configurable parameter set to match empirical observations in influenza (Bloom and Glassman, 2009; Rabadan et al., 2006)). Specifically, mutation events randomly select a nucleotide site and assign a new nucleotide by drawing from the K80-model probability distribution. We model purifying selection by rejecting a subset of mutations based on their properties. Mutations that introduce stop codons are rejected. Nonsynonymous mutations are probabilistically either accepted or rejected according to a user-defined “acceptance rate”, with separate acceptance rates for epitope and non-epitope sites. In this paper, we use acceptance rates of 0.75 and 0.25 at epitope and non-epitope sites, respectively, which we tuned to reproduce empirical H3N2 mutation statistics (Huddleston et al., 2020). The higher acceptance rate at epitope sites reflects positive selection for immune escape, whereas the lower rate at non-epitope sites reflects stronger purifying selection. Selection between hosts is a separate process in `antigen-prime`, and depends only on a virus’s antigenic coordinates and the immune memory of the host it contacts (eq. (2)).

The antigenic effect of a nucleotide mutation depends on if and how it changes the protein sequence (Figure 1B). Synonymous mutations do not alter the virus’s antigenic coordinates. Nonsynonymous mutations at epitope sites cause large movements in antigenic space with step sizes drawn from a gamma distribution with mean 0.6 antigenic units (analogous to the original antigen). Nonsynonymous mutations at non-epitope sites cause very small movements with step sizes drawn from a gamma distribution with mean 1 *×* 10*^−^*^4^ antigenic units. The direction of movement (*θ*) is random for both types of mutations. Thus, in `antigen-prime`, antigenic evolution is mostly driven by nonsynonymous mutations at epitope sites, while other mutations accumulate with littleto-no antigenic impact.

In summary, the main difference between antigen and `antigen-prime` is that virus objects in `antigen-prime` have sequences. In antigen, mutation events directly result in changes to the virus object’s location in antigenic space. In `antigen-prime`, mutation events first act on the virus’s sequence, which in turn can result in changes to the virus’s location in antigenic space. Other aspects of the simulator are the same, including antigenic selection on viruses to escape immune memory in the host population.

Related to antigenic selection, we have also updated `antigen-prime` to record population immunity over time. This feature enables benchmarking of variant-assignment methods by providing explicit fitness values for viruses. Every *t* days, the simulator samples *n* hosts and records each sampled host’s full immune history, that is, the antigenic coordinates of all of that host’s prior infections. These per-host immune histories enable fitness calculation for sampled viruses, where a virus’s fitness is its mean risk of infection across the sampled hosts.

### 2.2 Simulating long-term influenza evolution using **antigen**-prime

We used `antigen-prime` to simulate 30 years of H3N2 seasonal influenza evolution. We initialized simulations with the full-length HA sequence from A/Beijing/32/1992. Each simulation comprised three geographical demes: the Northern hemisphere, Southern hemisphere, and tropics, each with 30 million hosts. We ran 30 parallel replicate simulations to quantify variability in outcomes.

As with the original antigen, `antigen-prime` simulations reproduced influenza-like phylogenetic structure and rates of antigenic evolution. In natural H3N2 evolution, the average time between two contemporaneous sequences and their most recent common ancestor is expected to be *∼*3.22 years (Scotch et al., 2025), reflecting influenza’s spindly phylogenetic structure. The average rate of antigenic change is expected to be *∼*1.6 antigenic units per year, as quantified using experimental data from hemagglutination-inhibition assays (Hirst, 1943; Koel et al., 2013; Smith et al., 2004). Many `antigen-prime` simulations resulted in values close to these numbers (Figure S1).

`antigen-prime` simulations also reproduced influenza-like mutational patterns. As an empirical reference, we considered a phylogenetic tree that Huddleston et al. made using seasonal H3N2 influenza HA sequences collected over 25 years (Huddleston et al., 2020). For this tree, and for each of our simulated trees, we counted the number of epitope and non-epitope mutations observed along the branches of the tree, separately doing so for trunk branches and side branches, and using the set of epitope sites from Luksza and Lässig (Luksza and Lässig, 2014). Many of the simulated trees had counts similar to the empirical tree (Figure S2). Table 1 shows counts for a single simulation that reproduced realistic influenza-like dynamics in terms of mutational patterns, phylogenetic structure, and antigenic movement, and which we describe below in more detail. On the trunk, the simulated and empirical trees have similar mutation counts, and similar ratios in counts between epitope and non-epitope mutations. On side branches, the mutation counts are substantially higher for the simulated tree because this tree has substantially more sequences (this high sampling density was useful for downstream benchmarking purposes). However, the ratio of counts between epitope and non-epitope mutations, which is not sensitive to sampling density, is similar between the simulated and empirical trees. Given the complexity of natural influenza evolution and the simplifying assumptions in our model, we expect approximate rather than precise agreement with empirical values.

**Table 1:** Comparison of mutation patterns between an empirical phylogeny and the simulated phylogeny used for downstream analysis. The table shows counts of mutations occurring in epitope and non-epitope regions along the trunk vs. side branches. Empirical values are derived from 25 years of H3N2 HA sequences from Huddleston et al. (Huddleston et al., 2020), with trunk and side branch annotations following the methodology of Bedford et al. (Bedford et al., 2015). Scaled values extrapolate empirical counts to 30 years (*×*1.2) for comparison with simulation values. Simulation values are from a 30 year simulation.

| Mutation Characteristic | Empirical Value<br>(25 years) | Scaled Value<br>(30 years) | Simulation Value<br>(30 years) |
| --- | --- | --- | --- |
| *Trunk Mutations* |  |  |  |
| Epitope mutations | 50 | 60 | 50 |
| Non-epitope mutations | 32 | 38 | 26 |
| Ratio (epitope:non-epitope) | 1.56 | 1.56 | 1.92 |
| *Side Branch Mutations* |  |  |  |
| Epitope mutations | 485 | 582 | 959 |
| Non-epitope mutations | 1177 | 1412 | 3029 |
| Ratio (epitope:non-epitope) | 0.41 | 0.41 | 0.32 |

In the following sections, we benchmark methods for analyzing influenza sequences in detail on the single simulation summarized in Table 1 and Figure 2, and we assess how those results generalize across all 45 flu-like candidate simulations in the supplement. A phylogenetic tree of viruses sampled from this simulation exhibits a characteristic ladder-like structure (Figure 2A). Case counts across the three demes display realistic seasonal patterns, with Northern and Southern demes exhibiting opposite seasonal peaks and the tropics showing more constant transmission (Figure 2B). The distribution of sampled viruses in antigenic space reveals distinct antigenic clusters that show sustained antigenic evolution over time (Figure 2C). Epitope mutations accumulate approximately linearly over time at 0.83 mutations per year (*R*^2^ = 0.99), beginning from the roughly 14 mutations carried over from the discarded 10-year burn-in period (Figure 2D). Global variant frequency dynamics show sequential variant emergence-and-replacement patterns observed in the original `antigen` (Bedford et al., 2012) (Figure 2E). Panels A, B, C, and E confirm that `antigen-prime` recapitulates antigenic and epidemiological dynamics already demonstrated by the original antigen (Bedford et al., 2012), whereas panel D reflects a capability unique to `antigen-prime`: epitope mutations accumulating at a measurable rate that is tied to the underlying genetic sequence evolution.

**Figure 2:**
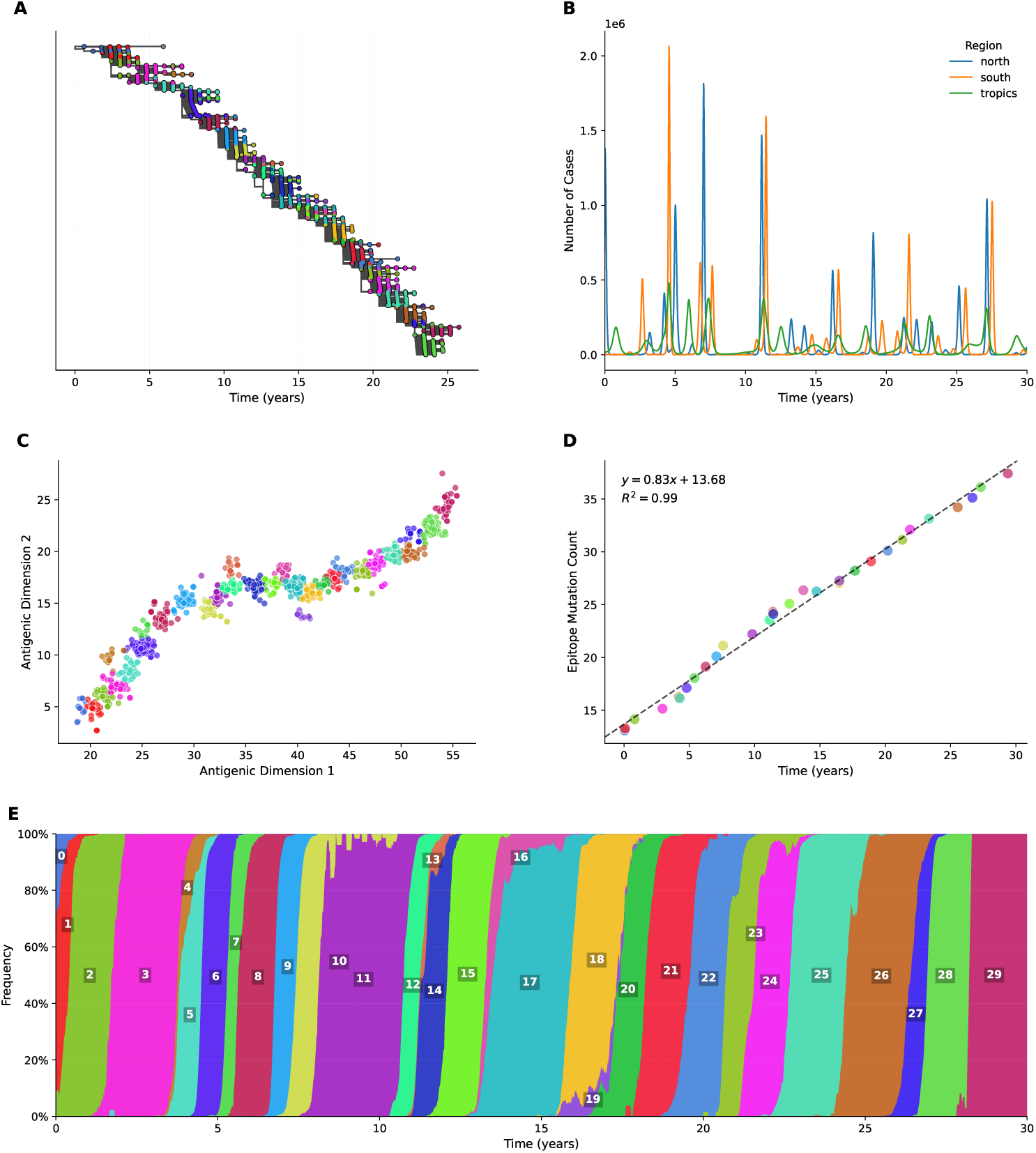
Genetic, antigenic, and epidemiological dynamics from a 30 year `antigen-prime` simulation. This is the single simulation that we carry through the benchmarks in the main text; performance across all 45 flu-like candidate simulations is reported in the supplement (Figure S4, Figure S5). **A)** Phylogenetic tree of 150,000 viruses sampled from the simulation, with tips colored by antigenic variant assignment. **B)** Case counts across three geographic demes (North, Tropics, South), demonstrating realistic seasonal epidemic patterns with hemispheric differences. **C)** Antigenic space of sampled viruses colored by variant, with variants assigned using k-means clustering on antigenic coordinates. The data show distinct antigenic clusters. **D)** Epitope mutations accumulate approximately linearly over time, indicating consistent antigenic drift throughout the simulation. The plotted 30-year window follows a discarded 10-year burn-in (see Methods), so approximately 14 epitope mutations have already accumulated at the start of the window. Each point is one antigenic variant, plotted at its mean sampling time and mean epitope mutation count; the black line is an ordinary least-squares fit to those points, with slope 0.83 mutations/year (*R*^2^ = 0.99). The fit is close to, though somewhat below, the rough expectation of one epitope mutation per year. **E)** Global variant frequency dynamics aggregated across all demes, showing sequential variant emergence and replacement patterns.

### 2.3 Benchmarking variant-assignment methods

We sought to use the above `antigen-prime` simulation to benchmark methods for grouping viral genetic sequences into variants with similar antigenicity, with the simulation providing ground-truth coordinates of each virus in antigenic space. We evaluated three methods. The first method uses k-means clustering of viruses by their ground-truth antigenic coordinates, providing a baseline for comparison. The second method uses Neher’s clade suggestion algorithm (Neher et al., 2025) to cluster viruses based on a phylogenetic tree built from their sequences. This approach defines variants by considering tree topology, overall genetic divergence, and amino acid changes at epitope sites, and has been applied in efforts to guide seasonal influenza vaccine composition (Huddleston et al., 2024). We parameterized this method using the same set of 49 epitope sites that we used in the simulation. The third method uses pathogen-embed (Nanduri et al., 2024) to compute pairwise Hamming distances between viral sequences, then use those data to project sequences into a lowdimensional space using t-SNE, and cluster sequences in that space using kmeans. We configured the antigenic and sequence-based methods to produce 30 variants, and the phylogenetic method to produce approximately 30 variants (resulting in 36) for comparison. When visualized in antigenic space, all three methods create largely distinct clusters (Figure 3A-C), even though the latter two approaches use only genetic data. As expected, the first approach results in strictly non-overlapping clusters. The latter two approaches result in clusters that are mostly separated, but have some overlap, reflecting the inherent difficulty in cleanly resolving antigenic boundaries from genetic data alone.

**Figure 3:**
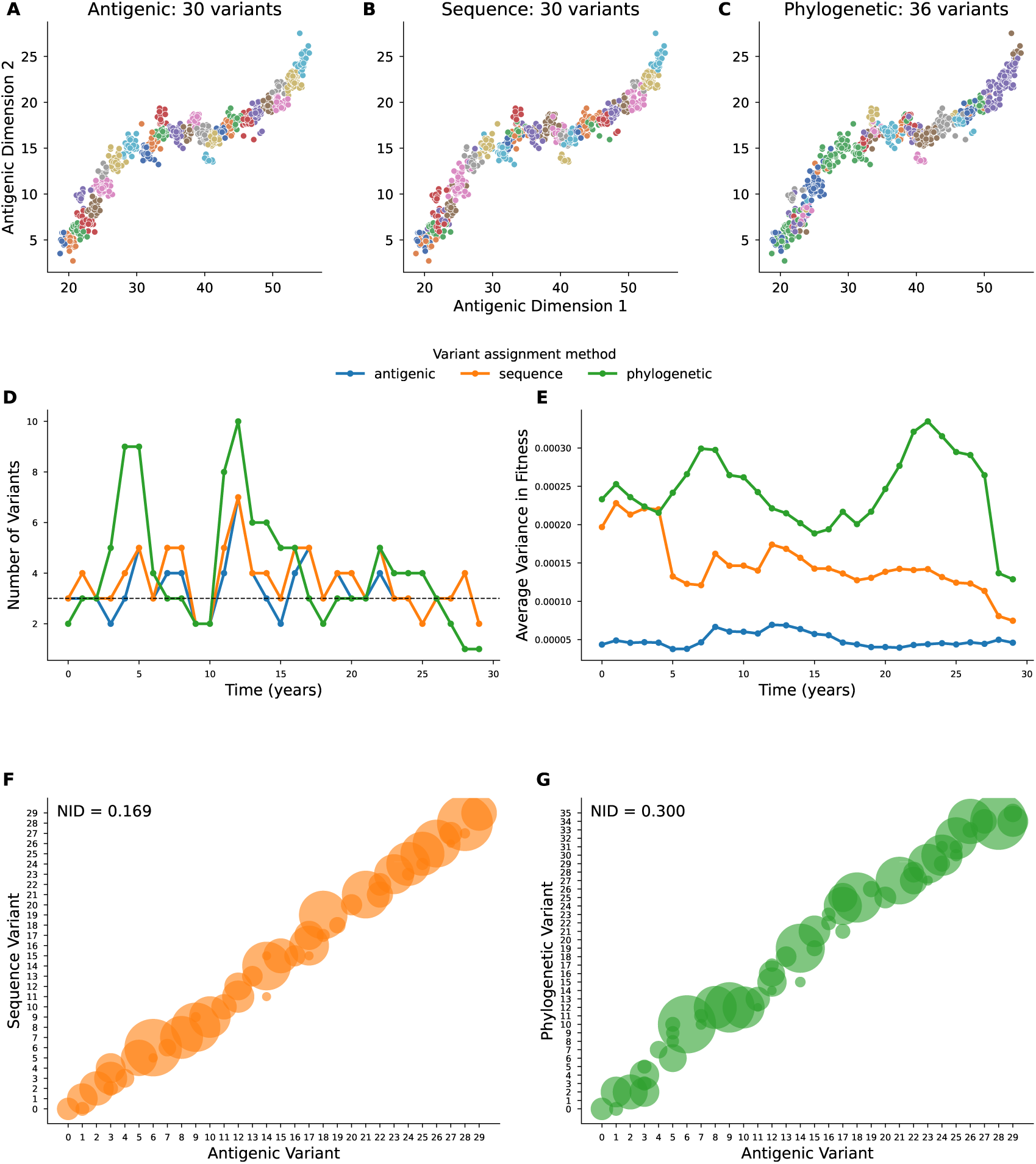
Benchmarking variant-assignment methods. **A-C)** Variant assignments visualized in simulated antigenic space. Each panel shows the same viral sequences plotted by their antigenic coordinates, with colors representing different variants. **D)** Number of variants circulating over time. The black dashed line marks three variants per year, the rate the *k* = 30 clustering parameter was chosen to produce over the 30-year simulation (see Methods). **E)** Average within-variant fitness variance over time. Lower values indicate better grouping of viruses with similar fitness. **F)** Variant-assignment agreement between the sequence-based method and groundtruth antigenic variants. Bubble sizes represent the number of sequences shared between compared variants. The overlap is high (NID = 0.169). **G)** Same as panel F, but for the phylogenetics-based method. The overlap is intermediate (NID = 0.300).

We quantified method performance using the following three metrics. First, we quantified the number of circulating variants per year. We tuned each method to produce approximately three per year to reflect real-world influenza dynamics (Huddleston et al., 2024). Each method resulted in similar traces of this quantity over time (Figure 3D).

Second, we quantified the variance in antigenic fitness among viruses assigned to a given variant (Figure 3E). Specifically, for each year, we calculated the fitness of each sampled virus from the entire simulation as its mean risk of infection across that year’s sampled host immune histories (eq. (2)), where a virus’s risk against a given host decreases with the antigenic distance to the nearest virus in that host’s immune memory. For each variant, we then computed the variance in these fitness values among viruses assigned to that variant, and then averaged these variances across all variants. The blue line from Figure 3E shows the performance of the first variant-assignment method, which clusters viruses by their ground-truth coordinates in antigenic space, and which we use as a baseline for evaluating the other two methods. The blue line is close to zero for all years, indicating low variance in fitness values and thus high performance. The other two methods also result in low variance (see the orange and green lines), as expected from their ability to effectively cluster viruses in antigenic space (Figure 3B/C). However, the variance of these methods is *∼*3-8fold higher than the blue baseline, reflecting the observation that there is some overlap between clusters. The sequence-based method shows consistently lower variance, and thus higher performance, than the phylogenetic method.

Third, we quantified the extent that the sequence- and phylogenetics-based assignments agreed with the ground-truth assignments from k-means clustering in antigenic space (Figure 3F,G). We quantified agreement using the normalized information distance (NID) (Li et al., 2004); a distance metric ranging from 0 to 1 where lower values indicate more agreement between two variant assignments and higher values indicate less agreement. Both the sequence-based and phylogenetic-based methods resulted in assignments with high overlap with ground-truth assignments, with the former method showing better overlap (NID = 0.169) than the latter (NID = 0.300).

Overall, this benchmark indicates that both the sequence-based and phylogenetic-based variant-assignment methods are effective at grouping viruses into antigenically similar variants, with the former method showing higher performance. These conclusions are consistent across all flu-like candidate simulations, not only the selected simulation (Figure S4). The success of these approaches also helps validate that `antigen-prime` successfully implements the fundamental coupling between genetic sequences and antigenic phenotypes that drives influenza evolution.

### 2.4 Benchmarking methods for inferring variant-specific growth rates

Next, we sought to use the simulated data to benchmark the ability of models to infer variant-specific growth rates. Natural data on SARS-CoV-2 has been used to benchmark the ability of MLR models to perform this task (Abousamra et al., 2024). But, previous studies have not benchmarked the more sophisticated FGA and GARW models.

To derive ground-truth growth rates from the simulated data, we divided the 30 years of data into overlapping one-year windows staggered every six months, capturing influenza seasons in both Northern and Southern demes. We further divided the simulated data by North, South, and Tropics demes. Then, for each window from each deme, we computed variant-specific frequencies and growth rates in weekly time bins, using variant assignments from k-means clustering of viruses in antigenic space. Specifically, we defined the empirical growth rate of variant *v* as the per-bin change in its log incidence,

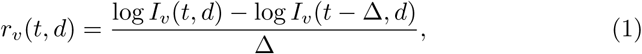

where *I_v_*(*t, d*) = *f_v_*(*t, d*) *C*(*t, d*) is the incidence of variant *v* in deme *d*, given by the product of its frequency *f_v_*(*t, d*) and the total case counts *C*(*t, d*), and Δ is the spacing between time bins in days. We omitted some variants at some time points due to insufficient case or sequence count data to accurately derive growth rates.

We then tested the ability of the FGA and GARW models from evofr to recover these growth rates. We separately fit each model to each window of data from each deme. As input, the models take the total number of reported cases among the host population and the observed counts for each variant in the sampled viral population. They then use a Bayesian approach to estimate probability distributions of variant-specific growth rates and frequencies over time. To analyze model predictions, we sampled 500 times from the inferred posterior distributions for variant growth rates and report the median values and 95 % highest posterior density (HPD) intervals. For each analysis window, we then calculated the mean absolute error (MAE) between predicted and groundtruth growth rates.

Below, we mostly focus on results from the GARW model as it allows growth advantages to vary smoothly over time, accommodating situations where the fixed-advantage assumption made in the FGA model may break down: for instance, due to shifting population immunity or cross-immunity between variants (Figgins and Bedford, 2025). Results for the simpler FGA model show comparable performance and are available in the supplement (Figure S5, Figure S6).

The performance of the GARW model varied across analysis windows. For many windows, the MAE values are low near zero, indicating accurate growthrate predictions (Figure 4). Figure 4B provides a detailed view of one such window, which shows good agreement between variant-specific frequencies and growth rates inferred by the model (see the lines) and those directly derived from the simulated data (see the circles).

**Figure 4:**
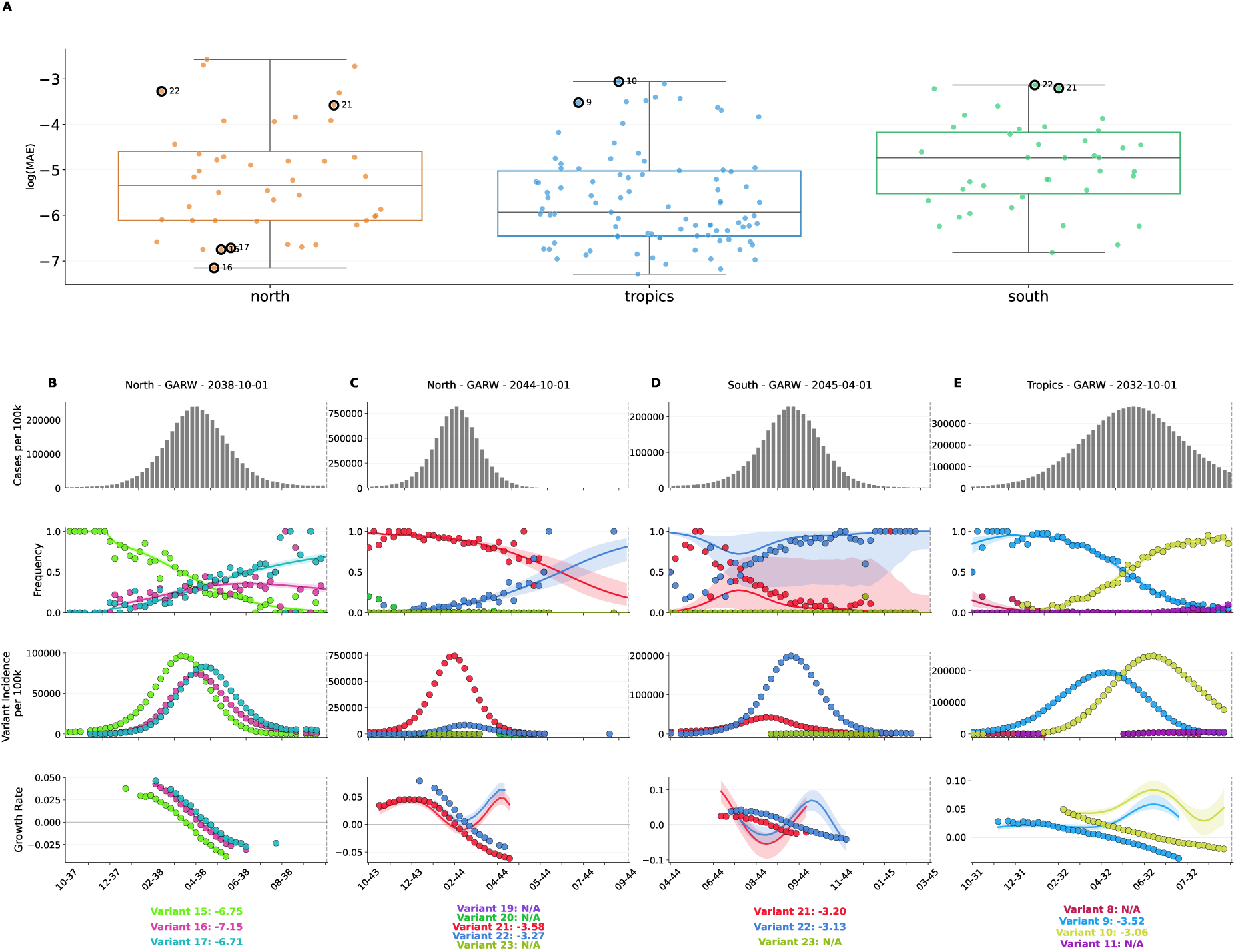
GARW model performance on variant growth-rate inference. **A)** Distribution of log MAE values of inferred variant-specific growth rates across geographic demes. Each data point represents a single variant from a specific training window. Variants selected for panels B-E are circled. **B-E)** Example analysis windows. Each panel shows case counts per 100,000 hosts over time (top), variant frequencies (second), case counts by variant (third), and inferred growth rates (bottom) for a given window. Growth rates reflect the rate of change of *absolute* variant incidence (variant frequency *×* total case counts; eq. (1)), so both variants can show negative growth rates during a seasonal decline in overall incidence even as the fitter variant rises in relative frequency. Points show values derived directly from the simulated data. Not all variants are shown at all time points due to data filtering on observed count thresholds for a series of timepoints (see Methods). Solid lines show median of model inferences and shaded regions indicate 95 % HPD intervals. Average log MAE values for each variant are reported below the growth-rate plots. **B)** Successful inference of frequencies and growth rates (North, 2038-10-01). **C)** Growth rates overestimated for variants 21 and 22 near the end of the window (North, 2044-10-01). **D)** Growth rates inaccurately inferred for variants 21 and 22 (South, 2045-04-01). **E)** Growth rates inferred as positive for variants 9 and 10 in time bins where they are observed to be negative (Tropics, 2032-10-01).

However, for several other windows, the MAE values are substantially higher, pointing to inaccurate growth-rate predictions. Examining these windows revealed a failure mode that has not been documented in previous studies. Figure 4C-E provides three examples of this failure mode. In each case, there is one variant that initially predominates and then begins to decline in frequency as one or two low-frequency variants start increasing in frequency. The model accurately predicts the frequency trajectory of each variant. However, in many weekly time bins, the model does not accurately predict variant-specific growth rates, especially in bins near the middle or end of the window when the initial predominant variant begins to substantially decline in frequency. Often, the prediction error is larger than the estimated model uncertainty (observed growth rates fall outside the HPD interval). Strikingly, there are multiple examples where the predicted and observed growth rates have opposing signs, like in Figure 4E where variants 9 and 10 are inferred to have positive growth rates in several time bins where they are actually negative, or in Figure 4C where the same occurs for variants 21 and 22. In general, in time bins with enough data to quantify growth rates for multiple variants, when the predicted growth rate is inaccurate for one variant, it tends to be inaccurate for the other variants by roughly the same amount and in roughly the same direction, indicating systematic bias. This bias may stem from an underlying assumption in both the GARW and FGA models that variant-specific growth rates tend to change in a concerted manner. To gauge how often such windows arise beyond the selected simulation, we scored every variant-window across the 45 flu-like candidate simulations and treated log(MAE) *> −*4 as elevated error (Figure S5). Of the 10,710 GARW variant-windows carrying enough data to score, 2,711 (25.3%) did not converge, and of the 7,999 that did, 1,085 (13.6%) exceeded that threshold. Elevated error is therefore not a rare event in these simulations. Inspecting flagged windows turned up more than one kind of pathology, and we describe the two we encountered most often rather than an exhaustive catalog. The first is a sharp decline in the number of circulating sequences or reported cases within the window, which leaves little data from which to resolve variant-specific rates. The second is a window in which overall incidence is falling while a fitter variant gains in relative frequency. There the model follows each variant’s frequency trajectory correctly but mis-estimates its growth rate, in several bins inferring a positive rate for a variant whose absolute incidence is in fact declining (Figure 4E). This is the same distinction between absolute incidence and relative frequency that eq. (1) makes explicit, and a variant gaining share during a seasonal trough is exactly where the two come apart.

In all, this benchmark indicates that the GARW and FGA models are largely effective at predicting variant-specific growth rates from the simulated data. However, it also revealed potential problems with these models that motivate additional investigation.

## 3 Discussion

We have presented `antigen-prime`, a simulator that jointly models the genetic and antigenic evolution of viruses under selection from host population immunity. We showed that `antigen-prime` can be used to simulate H3N2 seasonal influenza-like dynamics over long timescales. The dynamics are influenza-like in terms of their phylogenetic structure, antigenic evolution, and sequence-level mutation patterns, with mutations at epitope sites driving antigenic change. We then used the simulated data to benchmark computational methods that are currently used to interpret influenza surveillance data, and have relevance for public-health decision-making.

Our work helps address a fundamental challenge in evaluating these computational methods: the lack of ground-truth data in natural viral populations makes it difficult to assess whether the methods accurately capture the biological processes they aim to model. Simulated data with known ground-truth values can be used to benchmark methods. But, the field lacks simulators of pathogen evolution that model both genetic and antigenic evolution under selection from host population immunity. `antigen-prime` fills the gap, enabling benchmarking of the methods we analyzed in this paper.

The benchmarking results update our understanding of the efficacy of these methods. The benchmark on variant assignment showed that both the sequencebased and phylogenetic-based methods were effective at grouping viruses with similar antigenic properties even without explicit access to fitness or antigenic information. Interestingly, the sequence-based approach performed better than the phylogenetic one. The reason for this is not immediately evident to us, and could warrant future exploration. Future work could also benchmark variant assignment over shorter evolutionary time scales relevant for interpreting real-time influenza evolution. We note that performance on this benchmark is probably higher than expected for natural viral populations, since predicting phenotype from genotype is more challenging in nature due to factors not captured in `antigen-prime`, such epistasis between mutations and variable immunity in the host population.

The benchmark on growth-rate inference showed that the GARW and FGA methods performed well in most time windows, but also revealed windows where model predictions dramatically differed from ground-truth. While the GARW method is conceptually more flexible, this additional flexibility did not provide substantial advantages over the simpler FGA method in our benchmark. Investigating low-performance windows identified a previously undocumented failure mode. Thus, these results helped identify limitations to these methods, and could be used to help guide future method development. One caveat bounds how far they should be read: we ran both models as distributed, without model validation or hyperparameter selection, so what we characterize is the behavior a practitioner would get off the shelf rather than the best either model can achieve. Whether tuning would remove the failure modes we describe, or merely move them, is an open question. We focused this benchmark on the FGA and GARW models because they infer growth rates from variant frequencies together with case counts, the same quantities from which we derive our ground-truth growth rates, and because they reflect approaches currently used in influenza surveillance. We did not directly benchmark the fitness model of Luksza and Lässig (2014), despite its conceptual influence on `antigen-prime`, because that model predicts clade frequencies from a sequence-based fitness function rather than consuming case counts; adapting it to recover the case-count-based growth rates evaluated here would require re-deriving the model, which we leave to future work. In the future, `antigen-prime` simulated data could also be used to benchmark the ability of methods to forecast influenza evolution, which is also highly relevant to developing effective vaccines.

Despite capturing many features of viral evolution under immune selection, `antigen-prime` makes several simplifying assumptions that limit its biological realism. The fitness of simulated viruses is determined solely by their antigenic phenotype. However, other models of influenza evolution also model potential fitness costs of mutations at non-epitope sites (Luksza and Lässig, 2014), where mutations can disrupt HA’s ability to mediate viral entry. Additionally, the model maintains static epitope sites throughout the simulation, and employs a simple mutation model without epistatic interactions. The antigenic space itself is also a simplification. We represent antigenic phenotype in two dimensions, following the antigenic cartography of Smith et al. (2004), which showed that influenza antigenic relationships are largely captured in two dimensions. The original antigen model represents antigenic phenotype as an *N* -dimensional vector, and Bedford et al. (2012) found that population turnover proceeds primarily along a single antigenic axis, so little appears to be gained by adding dimensions. Higher-dimensional representations that might capture finer antigenic structure nonetheless remain a direction for future work. Because antigenic relationships are measured as Euclidean distances, cross-reactivity in our model is symmetric, so that immunity to one virus protects against a second exactly as much as the converse. Experimental measurements, for instance from hemagglutination-inhibition assays, show that this is not always the case in nature (Smith et al., 2004). We retain the Euclidean representation for its simplicity and because it is the conventional way of describing antigenic distances between influenza viruses; relaxing the symmetry assumption is another avenue for future work.

A further simplification concerns how a mutation’s antigenic effect is assigned. Because we drew the antigenic direction independently at each mutation event, a given amino-acid substitution does not carry a repeatable antigenic effect across genetic backgrounds, and the model is correspondingly not time-reversible: a lineage that acquires a substitution and later loses it is unlikely to return to its original antigenic position. A method built specifically to exploit convergence in antigenic phenotype, in which the same substitution is assumed to produce the same antigenic displacement wherever it arises, would therefore be looking for structure that our simulated data do not contain.

We do not, however, think this limitation substantially biases the benchmarks reported here. That is because convergent evolution at the sequence level is concentrated in rare lineages. Among more established lineages that variant assignment operates on, most epitope substitutions arise only once, and this concentration sharpens as the establishment threshold tightens (Figure S3D). On the occasions when an established substitution does recur, it tends to arise in genetically distant backgrounds and years apart (Figure S3E, F).

Our representation of host immunity is similarly simplified. Each host’s immunity is a set of discrete antigenic coordinates recorded from its prior infections, and susceptibility to a new virus is governed by the single nearest prior infection (the minimum antigenic distance; eq. (2)). This strategy ignores multiple factors that shape real host antibody responses (Fonville et al., 2014; Henry et al., 2021). Although `antigen-prime` can optionally model waning immunity by removing a randomly chosen entry from a host’s immune history at a constant per-host rate (the waning option), we disabled this feature for the present analysis, so hosts retain all prior infections indefinitely. The model also omits immune imprinting, whereby a host’s first childhood exposure shapes subsequent responses and creates systematic differences across birth cohorts (Arevalo et al., 2020; Cobey and Hensley, 2017; Gostic et al., 2016; Vieira et al., 2021). `antigen-prime` also omits vaccination, a dominant driver of population immunity in the settings the benchmarked methods target. A Cobey-lab fork of antigen does incorporate vaccination events (https://github.com/cobeylab/antigen-vaccine), though it does not carry the sequence model or the other extensions reported here, so the two capabilities are not currently available in a single simulator. This simplified representation is inherited from the antigen framework (Bedford et al., 2012) as a deliberate tradeoff toward tractability. Together with the antigenic-space assumptions above, these choices likely make the simulated immunity landscape smoother and simpler than what is observed in nature. Despite these simplifying assumptions, the simulated data still recapitulate multiple key aspects of influenza evolution that are relevant to the benchmarks; the failure of models to consistently predict variant growth rates even in this simplified scenario suggests they may also struggle to do so in more complex natural settings.

In all, `antigen-prime` provides a powerful framework for simulating seasonal influenza evolution, enabling researchers to benchmark and guide development of methods for interpreting influenza surveillance data. In the future, `antigen-prime` could be tuned to simulate evolutionary dynamics of other viruses, helping to benchmark methods for a variety of pathogens related to human health.

## 4 Methods

### 4.1 Data and code availability

The `antigen-prime` simulator source code is available at https://github.com/matsengrp/antigen-prime. Analysis scripts, simulation outputs, and code to generate all figures are available at https://github.com/matsengrp/antigen-forecasting.

### 4.2 Implementation of antigen-prime

`antigen-prime` is implemented as a Java program forked from the original antigen simulator (Bedford et al., 2012). The software is compiled using Maven and requires Java 11 or higher, with dependencies specified in the project’s pom.xml file. The complete source code is available at https://github.com/matsengrp/antigen-prime.

Two major extensions distinguish `antigen-prime` from the original simulator: (1) explicit genetic sequence evolution with site-specific mutation effects on antigenic movement, and (2) periodic sampling of host population immunity to enable fitness calculations for downstream analysis.

The core simulation algorithm follows the discrete-event SIR (SusceptibleInfected-Recovered) framework described in Bedford et al. (Bedford et al., 2012), with key extensions to couple genetic and antigenic evolution. The simulation proceeds through discrete time steps, modeling viral transmission dynamics across structured host populations (demes) while tracking both genetic sequences and antigenic coordinates for each virus. At each time step, the algorithm processes infection events based on host susceptibility and viral fitness, applies stochastic mutations to viral genomes according to the K80 model, updates antigenic coordinates based on mutation type and location, and samples host immune histories periodically to enable per-host fitness calculations for downstream analysis.

The risk that host *h* is infected upon contact with virus *v* is a piecewise-linear function of 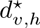, the minimum Euclidean antigenic distance between *v* and the antigenic coordinates stored in *h*’s immune memory:

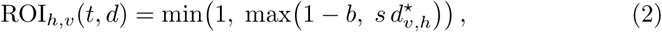

where *s* is the slope converting antigenic distance into infection risk (the smithConversion parameter) and *b* is the immunity conferred by an antigenically identical prior infection (the homologousImmunity parameter). A host with no relevant prior immunity (large 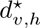) is therefore fully susceptible (ROI*_h,v_* = 1), whereas a host previously infected by an antigenically identical virus retains a residual infection risk of 1 *− b*. We used *s* = 0.07 and *b* = 0.95, matching the original antigen model (Bedford et al., 2012).

In this paper, we use an overall viral mutation rate of *µ* = 10*^−^*^3^ mutations per virus per day. Epitope sites comprise *∼*9% of the sequence (49/566 sites) and have an acceptance rate of 0.75, resulting in an effective mutation rate of *µ ×* 0.065 mutations per virus per day at epitope sites. Non-epitope sites comprise the remaining *∼*91% of the sequence and have a lower acceptance rate of 0.25, reflecting greater purifying selection at these sites, resulting in an effective rate of *µ ×* 0.228 mutations per virus per day. We tuned these acceptance rates to reproduce mutational patterns observed in seasonal influenza in nature (Figure S2).

### 4.3 Simulation parameterization and host population immunity sampling

We simulated 40 years of influenza evolution across three demes, and discarded the first 10 years as burn-in. We parameterized mutation step-size distributions to reflect the differential antigenic impact of epitope versus non-epitope mutations. Non-epitope sites used *r_ne_ ∼* Gamma(*α* = 1*, β* = 0.0001) while epitope sites used *r_e_ ∼* Gamma(*α* = 2.25*, β* = 0.267). All mutations received a step direction *θ ∼* Uniform(0, 2*π*). Because the step direction is drawn independently for each mutation event, the same amino-acid substitution generally produces different antigenic displacements in different sequence backgrounds rather than a single, repeatable effect. `antigen-prime` can instead assign fixed, pre-defined antigenic vectors to specific amino-acid changes, which make such a substitution produce a consistent displacement wherever it arises; we disabled this option here because it interfered with sustained antigenic evolution over the multi-decade simulations.

We sampled 150,000 viruses over the course of the simulation, proportionally by prevalence across demes and time, yielding approximately 7,606 unique nucleotide sequences to provide adequate count data for downstream growth-rate inference. To track population-immunity dynamics, we sampled 10,000 hosts from each deme every 365 days and saved each sampled host’s full immune history, that is, the antigenic coordinates of all viruses stored in that host’s immune memory. We use these per-host immune histories to compute virus fitness values in the downstream variant-assignment benchmark. The fitness of virus *v* at time *t* is its mean risk of infection across the hosts sampled at that time:

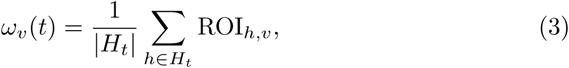

where *H_t_* is the set of hosts sampled at time *t*, pooled across demes, and ROI*_h,v_* is the risk of infection of host *h* by virus *v* (eq. (2)). Hosts with no prior immunity contribute the maximum risk of 1.0, since an empty immune memory leaves them fully susceptible.

### 4.4 Simulation selection for benchmarking applications

We applied three criteria for selecting simulations suitable for benchmarking: (1) realistic epidemiological dynamics with seasonal epidemic patterns in the temperate demes and year-round transmission in the tropics, (2) summary statistics matching empirical A/H3N2 HA genetic and antigenic evolution, and (3) sufficient viral diversity for robust benchmarking of methods for variant assignment and growth-rate inference.

To achieve the first criterion, we maintained the same antigenic and epidemiological parameter values from the original antigen paper by Bedford *et al*. (Bedford et al., 2012). We also set the overall mutation rate to *µ* = 10*^−^*^3^ mutation events per individual per day. For the second criterion, we ran 120 total simulations with four different parameter configurations for epitope and non-epitope mutation acceptance rates (epitope: 0.75 or 1.0; non-epitope: 0.2 or 0.25), using 30 replicates for each configuration. Of these, 85 ran to completion and produced the full set of genealogical and mutational summary statistics, and none of the completed simulations underwent viral population extinction (Figure S1, Figure S2). The remaining 35 did not run to completion and are excluded from all analyses reported here.

We define the mean pairwise genealogical diversity *π_G_* as the average total branch length between randomly sampled virus pairs:

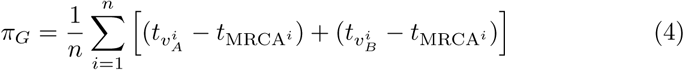

where *n* is the number of sampled pairs, *t_v_* is the birth time of virus *v*, and *t*_MRCA_ is the birth time of the most recent common ancestor for each pair. We then applied a filtering criterion to focus on “flu-like” simulations: *π_G_ ≤* 9.0 years, the threshold used by Bedford *et al*. (Bedford et al., 2012). All 85 completed simulations satisfied this criterion, spanning *π_G_* of 2.8 to 5.5 years, so it excluded none of them; the reduction from 120 to 85 reflects the simulations that did not run to completion rather than any filtering on diversity. Applying the remaining criteria, which required influenza-like phylogenetic structure, antigenic movement, and mutational patterns, identified 45 flu-like candidate simulations, and it is these 45 that we refer to as candidates throughout. For downstream variant assignment and growth rate inference benchmarking, we selected a single simulation with epitope acceptance rate of 0.75 and non-epitope acceptance rate of 0.25. This simulation had a simulated *π_G_* of 3.4 years and TMRCA of 3.3 years, and average antigenic movement of 1.5 units per year, closely matching empirical observations. The mutation summary statistics reported in Table 1 represent this selected simulation. Final selection involved confirming that phylogenetic trees and antigenic space distributions exhibited realistic influenza-like dynamics by visual inspection. Phylogenetic trees were inspected to ensure they displayed ladder-like structures, and case count dynamics were examined to confirm seasonal epidemic patterns in temperate demes and year-round transmission in the tropics.

### 4.5 Variant assignment for antigen-prime simulations

Three variant-assignment methods were applied to the 7,606 unique sequences: antigenic clustering (ground truth), sequence-based clustering, and phylogenetic variant assignment. Antigenic variants were defined by k-means clustering (*k* = 30) on two-dimensional antigenic coordinates. Sequence-based variants were assigned using pathogen-embed (Nanduri et al., 2024): (1) sequence alignment with MAFFT via augur align, (2) pairwise Hamming distance calculation, (3) t-SNE embedding in 2D space, and (4) k-means clustering (*k* = 30) on embeddings. The *k* = 30 parameter for both methods reflects empirical influenza dynamics of approximately three variants per year over the 30 year simulation.

Phylogenetic variants were assigned using the Neher clade assignment algorithm (Neher et al., 2025) following phylogeny reconstruction and ancestral inference. The algorithm was configured to use the same epitope sites defined by Luksza and Lässig (Luksza and Lässig, 2014) that are used in the `antigen-prime` simulation. Phylogeny inference used IQ-TREE via augur tree with augur refine refinement. Ancestral reconstruction applied augur ancestral and augur translate with default parameters. Clade assignment used parameters: bushiness branch scale 1.0, divergence scale 2.0, branch length scale 2.0, minimum clade size 22 sequences, targeting approximately 30 variants (resulting in 36). All variant assignments were re-labeled chronologically by average birth date of constituent viruses.

Within-variant fitness variance was calculated to evaluate how well each method grouped viruses with similar fitness. We computed the fitness of all viruses annually (eq. (3)), then calculated the average within-variant fitness variance for each assignment method.

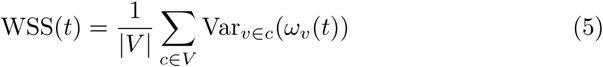

where *V* is the set of all variants assigned by a method, *|V |* is the total number of variants, *c* denotes a single variant (a set of viruses), and Var*_v∈c_*(*ω_v_*(*t*)) is the variance of the fitness values *ω_v_*(*t*) (eq. (3)) over the viruses *v* assigned to variant *c* at time *t*.

Agreement between variant assignments was quantified using the normalized information distance (NID) (Li et al., 2004):

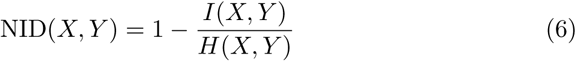

where *I*(*X, Y*) is the mutual information between assignments *X* and *Y*, and *H*(*X, Y*) is their joint entropy. NID ranges from 0 (identical assignments) to 1 (completely independent assignments).

### 4.6 Inferring variant growth rates with evofr forecasting models

We implemented two forecasting models from the evofr (Figgins and Bedford, 2025) framework to infer variant growth rates from simulated surveillance data. The Fixed Growth Advantage (FGA) model implements a renewal equation approach where each variant has a fixed multiplicative growth factor, while the Growth Advantage Random Walk (GARW) model allows variant growth advantages to vary smoothly over time using a random walk prior.

Data preparation involved extracting weekly variant-specific sequence counts and case counts from simulation outputs for each analysis timepoint. Data were separated by deme with analysis dates representing both in-season and out-of-season periods across different epidemic contexts. The retrospective observation window was limited to 365 days from each analysis date.

Model fitting used the evofr software package for both FGA and GARW models. Model-specific hyperparameters were initialized with spline basis functions of order 4 with 10 knots to model time-varying parameters. Generation time and reporting delay distributions were defined for the renewal equation models, with generation times parameterized as gamma distributions: *g*(*τ*) *∼* Gamma(mean = 3.0, std = 1.2). Sequence count data used Dirichlet-Multinomial likelihood with concentration parameter 100, while case count data used Negative Binomial likelihood with dispersion parameter 0.05. Variational inference approximated posterior distributions of model parameters using full-rank variational inference with 50,000 iterations and learning rate of 0.01, generating 500 posterior samples per model. We applied these settings as provided rather than selecting them for this benchmark, and carried out no model validation and no hyperparameter search. We focused on the inferred variant growth rates in this work.

### 4.7 Benchmarking growth rate inference performance

Empirical growth rates (*r*_data_*_,v_*) were calculated from simulated surveillance data using spline-based smoothing to reduce noise. For each variant *v* and location *d*, sequence data processing used univariate spline interpolation by log-transforming sequence counts to handle data skew and stabilize variance, applying a cubic univariate spline (degree *k* = 3) with smoothing factor *s* = 1.0 to the log-transformed data, then transforming the smoothed values back to the original scale.

After obtaining smoothed sequence counts, we calculated variant-specific incidence by multiplying total case counts by variant frequencies, then computed empirical growth rates as the change in log-transformed variant incidence over time:

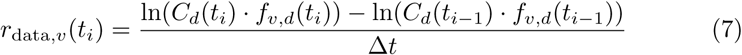

where *C_d_*(*t_i_*) represents the total case counts in location *d* at time *t_i_*, *f_v,d_*(*t_i_*) represents the smoothed frequency of variant *v* in location *d* at time *t_i_*, and Δ*t* is the time difference between observations. This approach scales the total disease burden by variant-specific frequencies, providing a representation of variant-specific growth rates.

Data filtering ensured reliable growth-rate estimates by excluding time points with smoothed sequence counts below 10 sequences (due to high sampling variance), variant frequencies below 1% of the total population (to avoid stochastic effects in rare variants), and variant incidence below 50 cases on any given day (to ensure sufficient case data). We required a minimum of 3 consecutive valid time points for a variant to be included in the growth-rate benchmarking analysis.

Performance on growth-rate inference was evaluated using mean absolute error (MAE) between the medians of the inferred growth rate posteriors (*r*_model_*_,v_*) and empirical (*r*_data*,v*_) growth rates for each variant:

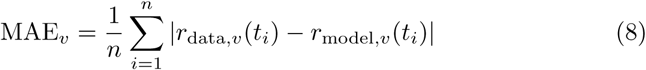

where *n* is the number of time points for variant *v*. We report log(MAE) for each variant to facilitate comparison across the small error ranges typically observed. To identify analysis windows with unforeseen pathologies, we flagged variant-windows whose per-variant growth-rate error exceeded log(MAE) = *−*4, restricting to converged fits with at least three scored time points, and then examined the flagged windows individually. Complete results, including detailed performance metrics for all analysis windows, are available at https://github.com/matsengrp/antigen-forecasting/notebooks/.

---

**Figure S1:**
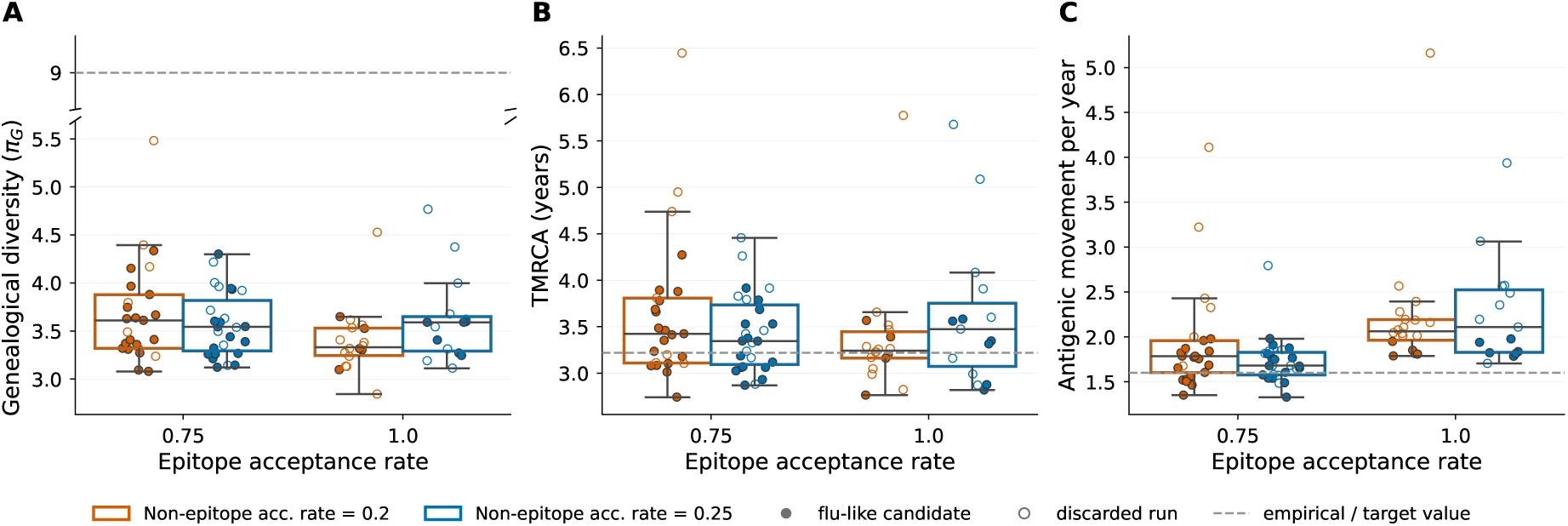
Genealogical and antigenic summary statistics for 30 years of H3N2-like evolution. Each point represents a single simulation; 85 of the 120 simulations are shown, the remaining 35 having failed to run to completion within their allocated resource time. Filled points mark the flu-like candidate simulations carried forward to the benchmarking analyses, and open points the remainder. Gray dashed horizontal lines represent empirical values reported in previous studies, and where such a value falls far outside the simulated range the axis is broken so the data keep their own scale. The 9.0 diversity cutoff was used in Bedford et al. (Bedford et al., 2012), and the antigenic movement per year metric was chosen to reflect results observed in Smith et al. (Smith et al., 2004) and Koel et al. (Koel et al., 2013). A) Genealogical diversity (*π_G_*). B) Time to most recent common ancestor (TMRCA), dashed line marks a target value based on data from nature (Scotch et al., 2025). C) Antigenic movement per year, dashed line marks a target value based on data from nature.

**Figure S2:**
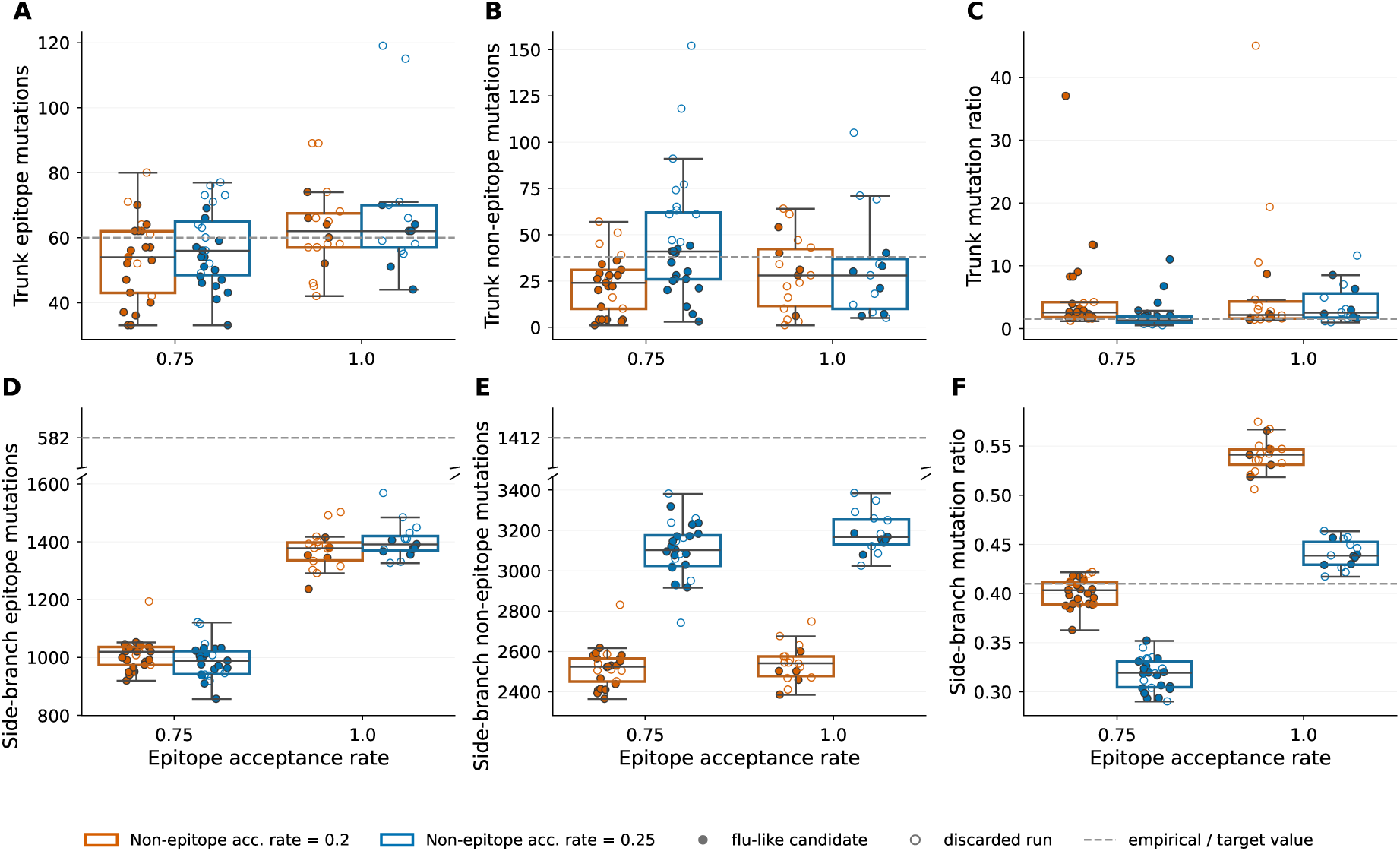
Summary of `antigen-prime` mutation statistics for 30 years of evolution across various epitope/non-epitope acceptance rate configurations. Each point represents results from a single simulation; 85 of the 120 simulations are shown, the remaining 35 having failed to run to completion within their allocated resource time. Filled points mark the flu-like candidate simulations carried forward to the benchmarking analyses, and open points the remainder. Gray dashed horizontal lines represent empirical values reported in Table 1, and where such a value falls far outside the simulated range the axis is broken so the data keep their own scale. **A)** Total number of epitope mutations observed on the trunk of the phylogeny. **B)** Total number of non-epitope mutations observed on the trunk of the phylogeny. **C)** Ratio of epitope to non-epitope mutations observed on the trunk of the phylogeny. **D)** Total number of epitope mutations observed on side branches of the phylogeny. **E)** Total number of non-epitope mutations observed on side branches of the phylogeny. **F)** Ratio of epitope to non-epitope mutations observed on side branches of the phylogeny.

**Figure S3:**
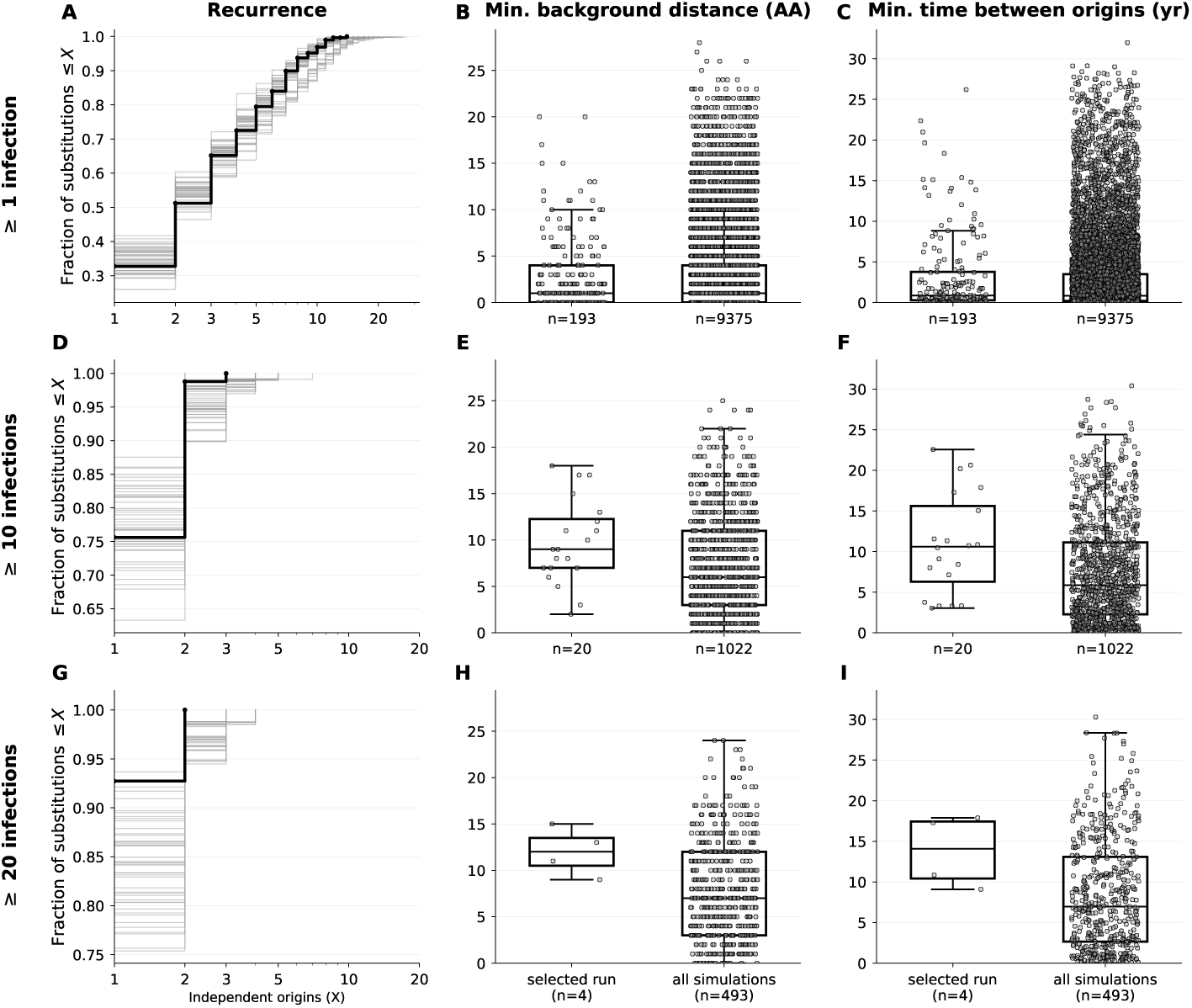
Recurrence of epitope substitutions among established lineages is rare, and grows rarer as the establishment threshold tightens, while the independent origins of the substitutions that do recur tend to arise on genetically and temporally distant backgrounds, across the 45 flu-like candidate `antigen-prime` simulations. Every quantity is computed from the true simulation genealogy, in which each branch carries the exact parent and child sequence, so the independent origins of a substitution are counted exactly rather than estimated from a reconstructed tree. The three rows restrict origins to those whose clade retains at least one (A–C), ten (D–F), or twenty (G–I) sampled infections; the middle row uses the threshold applied elsewhere, since variant assignment operates on lineages that spread. **A)** Cumulative fraction of epitope substitutions arising on at most *X* separate branches, so the value at *X* = 1 is the fraction arising only once; each faint line is one simulation and the selected simulation is bold, with the corresponding curves for the stricter thresholds in D and G. The curve lifts toward *X* = 1 as the threshold tightens, so recurrence is progressively confined to the transient lineages that the stricter cutoffs exclude. **B)** Minimum amino-acid distance between the genetic backgrounds of any two independent origins of a recurrent epitope substitution, one point per substitution, shown for the selected simulation and pooled across all candidates, with the stricter thresholds in E and H. **C)** Minimum time between any two independent origins of a recurrent epitope substitution, on the same terms, with the stricter thresholds in F and I. Both spreads widen with the threshold, so among the lineages that spread widely the independent origins of a recurrent substitution are separated by more amino acids and more years.

**Figure S4:**
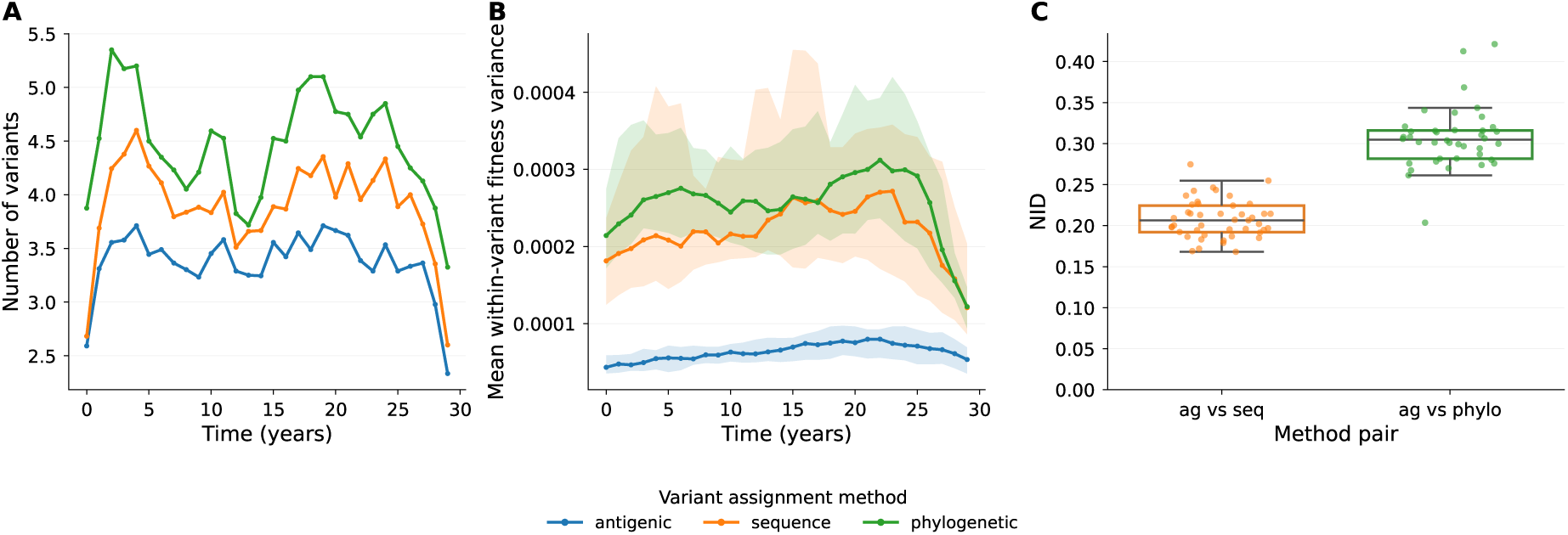
Variant-assignment performance pooled across all flu-like candidate simulations. This is the across-simulation counterpart to Figure 3, showing that the relative performance of the three variant-assignment methods is consistent across simulations rather than specific to the selected simulation. In panels A and B, lines show the mean across candidate simulations and shaded bands show the interquartile range. **A)** Number of variants circulating over time for each method. **B)** Mean within-variant fitness variance over time for each method; lower values indicate better grouping of viruses with similar fitness. **C)** Variant-assignment agreement, quantified by the normalized information distance (NID) between the ground-truth antigenic assignment and the sequence-based and phylogenetic assignments, with each point a single candidate simulation.

**Figure S5:**
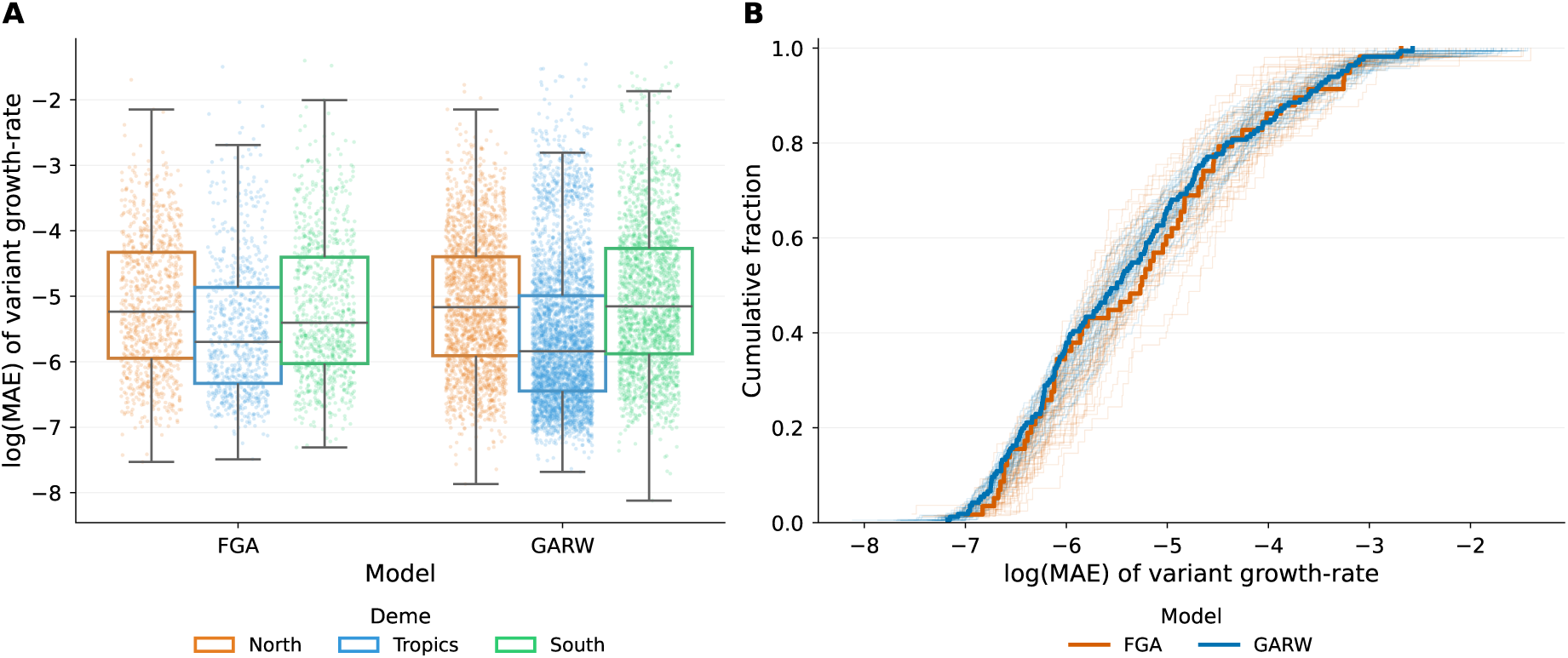
Variant-specific growth-rate inference error for the FGA and GARW models, pooled across all flu-like candidate simulations. **A)** Distribution of log(MAE) of inferred variant growth-rates by model (FGA, GARW) and geographic deme, across all candidate simulations. Both models show comparable performance across demes. **B)** Empirical cumulative distribution of log(MAE) for each candidate simulation (faint curves, colored by model); the bold curves are the selected simulation used in the main text. The analysis windows examined in Figure S6 were selected from the upper tail of this distribution.

**Figure S6:**
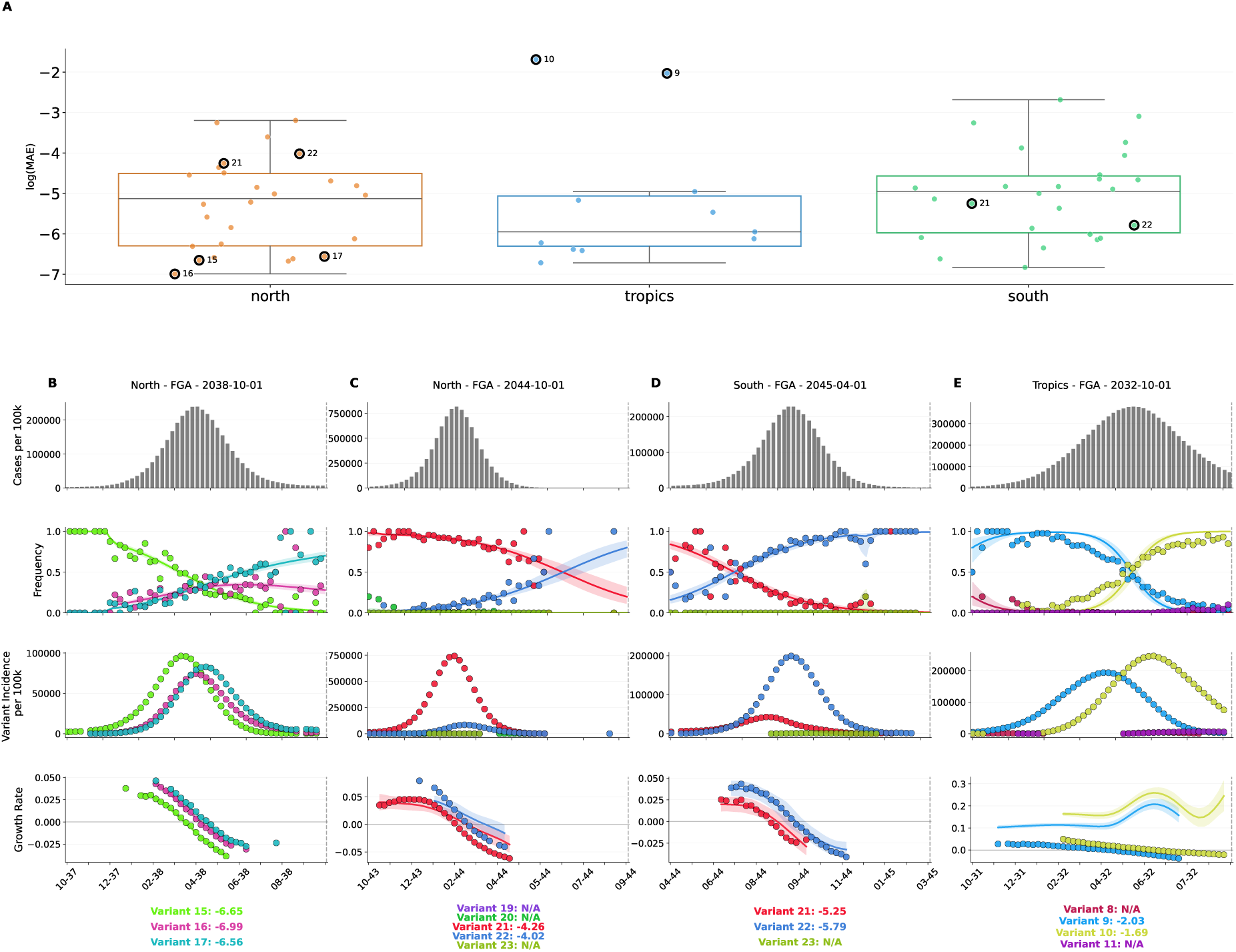
FGA model performance for variant growth-rate inference. **A)** Distribution of log MAE values of inferred variant-specific growth rates across geographic demes. Each data point represents a single variant from a specific training window. Variants selected for panels B-E are circled. **B-E)** Examples of FGA model performance across different training windows. Average log MAE values for each variant are reported below the growth-rate plots. **B)** Successful inference of frequencies and growth rates (North, 2038-10-01). **C)** Growth rates inaccurately inferred for variants 21 and 22 (North, 2044-10-01). **D)** Accurate inference of frequencies and growth rates (South, 2045-04-01). **E)** A non-converged FGA fit for variants 9 and 10, whose frequency nowcast appears accurate while the inferred growth rates are unreliable (Tropics, 2032-10-01). The two high-error tropics points circled in panel A correspond to this non-converged window, included to show that a fit whose frequencies look accurate can still yield unreliable growth-rate inferences.

## References

E. Abousamra, M. Figgins, and T. Bedford. Fitness models provide accurate short-term forecasts of SARS-CoV-2 variant frequency. PLOS Computational Biology, 20(9):1–20, 09 2024. doi: 10.1371/journal.pcbi.1012443. URL https://doi.org/10.1371/journal.pcbi.1012443.

P. Arevalo, H. Q. McLean, E. A. Belongia, and S. Cobey. Earliest infections predict the age distribution of seasonal influenza A cases. eLife, 9:e50060, 2020. doi: 10.7554/eLife.50060.

T. Bedford, A. Rambaut, and M. Pascual. Canalization of the evolutionary trajectory of the human influenza virus. BMC Biol., 10:38, Apr. 2012.

T. Bedford, S. Riley, I. G. Barr, S. Broor, M. Chadha, N. J. Cox, R. S. Daniels, C. P. Gunasekaran, A. C. Hurt, A. Kelso, A. Klimov, N. S. Lewis, X. Li, J. W. McCauley, T. Odagiri, V. Potdar, A. Rambaut, Y. Shu, E. Skepner, D. J. Smith, M. A. Suchard, M. Tashiro, D. Wang, X. Xu, P. Lemey, and C. A. Russell. Global circulation patterns of seasonal influenza viruses vary with antigenic drift. Nature, 523(7559):217–220, July 2015.

J. D. Bloom and M. J. Glassman. Inferring stabilizing mutations from protein phylogenies: application to influenza hemagglutinin. PLoS Comput. Biol., 5 (4):e1000349, Apr. 2009.

S. Cobey and S. E. Hensley. Immune history and influenza virus susceptibility. Current Opinion in Virology, 22:105–111, 2017. doi: 10.1016/j.coviro.2016.12.004.

M. D. Figgins and T. Bedford. Inferring variant-specific effective reproduction numbers from combined case and sequencing data. Sept. 2025. doi: 10.7554/elife.104802.1. URL https://doi.org/10.7554/eLife.104802.1.

J. M. Fonville, S. H. Wilks, S. L. James, A. Fox, M. Ventresca, M. Aban, L. Xue, T. C. Jones, N. M. H. Le, Q. M. Pham, et al. Antibody landscapes after influenza virus infection or vaccination. Science, 346(6212):996–1000, 2014. doi: 10.1126/science.1256427.

K. M. Gostic, M. Ambrose, M. Worobey, and J. O. Lloyd-Smith. Potent protection against H5N1 and H7N9 influenza via childhood hemagglutinin imprinting. Science, 354(6313):722–726, 2016. doi: 10.1126/science.aag1322.

J. Hadfield, C. Megill, S. M. Bell, J. Huddleston, B. Potter, C. Callender, P. Sagulenko, T. Bedford, and R. A. Neher. Nextstrain: real-time tracking of pathogen evolution. Bioinformatics, 34(23):4121–4123, Dec. 2018.

C. Henry, N.-Y. Zheng, M. Huang, A. Cabanov, K. T. Rojas, K. Kaur, S. F. Andrews, A.-K. E. Palm, Y.-Q. Chen, Y. Li, et al. Preexisting immunity shapes distinct antibody landscapes after influenza virus infection and vaccination in humans. Science Translational Medicine, 13(573):eabd3601, 2021. doi: 10.1126/scitranslmed.abd3601.

G. K. Hirst. Studies of antigenic differences among strains of influenza a by means of red cell agglutination. J. Exp. Med., 78(5):407–423, Nov. 1943.

J. Huddleston, J. R. Barnes, T. Rowe, X. Xu, R. Kondor, D. E. Wentworth, L. Whittaker, B. Ermetal, R. S. Daniels, J. W. McCauley, S. Fujisaki, K. Nakamura, N. Kishida, S. Watanabe, H. Hasegawa, I. Barr, K. Subbarao, P. Barrat-Charlaix, R. A. Neher, and T. Bedford. Integrating genotypes and phenotypes improves long-term forecasts of seasonal influenza A/H3N2 evolution. Elife, 9, Sept. 2020.

J. Huddleston, T. Bedford, J. Chang, J. Lee, and R. A. Neher. Seasonal influenza circulation patterns and projections for february 2024 to february 2025. 2024.

K. Ito, C. Piantham, and H. Nishiura. Predicted dominance of variant Delta of SARS-CoV-2 before Tokyo Olympic Games, Japan, July 2021. Euro Surveill., 26(27), July 2021.

A. Jariani, C. Warth, K. Deforche, P. Libin, A. J. Drummond, A. Rambaut, F. A. Matsen, Iv, and K. Theys. SANTA-SIM: simulating viral sequence evolution dynamics under selection and recombination. Virus Evol, 5(1): vez003, Jan. 2019.

C. Kikawa, A. N. Loes, J. Huddleston, M. D. Figgins, P. Steinberg, T. Griffiths, E. M. Drapeau, H. Peck, I. G. Barr, J. A. Englund, S. E. Hensley, T. Bedford, and J. D. Bloom. High-throughput neutralization measurements correlate strongly with evolutionary success of human influenza strains. eLife, Nov. 2025. doi: 10.7554/elife.106811.2. URL https://doi.org/10.7554/eLife.106811.2.

M. Kimura. A simple method for estimating evolutionary rates of base substitutions through comparative studies of nucleotide sequences. J. Mol. Evol., 16(2):111–120, Dec. 1980.

B. F. Koel, D. F. Burke, T. M. Bestebroer, S. van der Vliet, G. C. M. Zondag, G. Vervaet, E. Skepner, N. S. Lewis, M. I. J. Spronken, C. A. Russell, M. Y. Eropkin, A. C. Hurt, I. G. Barr, J. C. de Jong, G. F. Rimmelzwaan, A. D. M. E. Osterhaus, R. A. M. Fouchier, and D. J. Smith. Substitutions near the receptor binding site determine major antigenic change during influenza virus evolution. Science, 342(6161):976–979, Nov. 2013.

F. Krammer, G. J. D. Smith, R. A. M. Fouchier, M. Peiris, K. Kedzierska, P. C. Doherty, P. Palese, M. L. Shaw, J. Treanor, R. G. Webster, and A. GarćıaSastre. Influenza. Nat. Rev. Dis. Primers, 4(1):3, June 2018.

M. Li, X. Chen, X. Li, B. Ma, and P. M. B. Vitanyi. The similarity metric. IEEE Trans. Inf. Theory, 50(12):3250–3264, Dec. 2004.

M. Luksza and M. Lässig. A predictive fitness model for influenza. Nature, 507 (7490):57–61, Mar. 2014.

D. H. Morris, K. M. Gostic, S. Pompei, T. Bedford, M. Luksza, R. A. Neher, B. T. Grenfell, M. Lässig, and J. W. McCauley. Predictive modeling of influenza shows the promise of applied evolutionary biology. Trends Microbiol., 26(2):102–118, Feb. 2018.

N. Moshiri, M. Ragonnet-Cronin, J. O. Wertheim, and S. Mirarab. FAVITES: simultaneous simulation of transmission networks, phylogenetic trees and sequences. Bioinformatics, 35(11):1852–1861, June 2019.

S. Nanduri, A. Black, T. Bedford, and J. Huddleston. Dimensionality reduction distills complex evolutionary relationships in seasonal influenza and SARSCoV-2. Virus Evolution, 10(1):veae087, 11 2024. ISSN 2057-1577. doi: 10.1093/ve/veae087. URL https://doi.org/10.1093/ve/veae087.

R. A. Neher, J. Huddleston, T. Bedford, N. S. Lewis, R. Harvey, M. Galiano, A. M. P. Byrne, S. James, D. Smith, M. Luksza, D. Ruchnewitz, M. Laessig, S. Fujisaki, S. Watanabe, H. Hasegawa, N. Hassell, D. E. Wentworth, R. Kondor, Y. M. Deng, C. Dapat, K. Subbarao, and I. Barr. Nomenclature for tracking of genetic variation of seasonal influenza viruses. medRxiv, page 2025.12.06.25341755, Dec. 2025.

F. Obermeyer, M. Jankowiak, N. Barkas, S. F. Schaffner, J. D. Pyle, L. Yurkovetskiy, M. Bosso, D. J. Park, M. Babadi, B. L. MacInnis, J. Luban, P. C. Sabeti, and J. E. Lemieux. Analysis of 6.4 million SARS-CoV-2 genomes identifies mutations associated with fitness. Science, 376(6599):1327–1332, June 2022.

N. Ochsner, J. Bouman, T. G. Vaughan, T. Stadler, S. Bonhoeffer, and R. R. Regoes. Viral simulation reveals overestimation bias in withinhost phylodynamic migration rate estimates under selection. bioRxiv, page 2025.01.06.631458, Jan. 2025.

V. N. Petrova and C. A. Russell. The evolution of seasonal influenza viruses. Nat. Rev. Microbiol., 16(1):47–60, Jan. 2018.

C. Piantham, N. M. Linton, H. Nishiura, and K. Ito. Estimating the elevated transmissibility of the B.1.1.7 strain over previously circulating strains in England using GISAID sequence frequencies. bioRxiv, page 2021.03.17.21253775, Mar. 2021.

R. Rabadan, A. J. Levine, and H. Robins. Comparison of avian and human influenza a viruses reveals a mutational bias on the viral genomes. J. Virol., 80(23):11887–11891, Dec. 2006.

M. Scotch, T. O. C. Faleye, J. M. Wright, S. Finnerty, R. U. Halden, and A. Varsani. Campus-based genomic surveillance uncovers early emergence of a future dominant a(H3N2) influenza clade. iScience, 28(12):113941, Dec. 2025.

Y. Shu and J. McCauley. GISAID: Global initiative on sharing all influenza data from vision to reality. Euro Surveill., 22(13):30494, Mar. 2017.

D. J. Smith, A. S. Lapedes, J. C. de Jong, T. M. Bestebroer, G. F. Rimmelzwaan, A. D. M. E. Osterhaus, and R. A. M. Fouchier. Mapping the antigenic and genetic evolution of influenza virus. Science, 305(5682):371–376, July 2004.

M. C. Vieira, C. M. Donato, P. Arevalo, G. F. Rimmelzwaan, T. Wood, L. Lopez, Q. S. Huang, V. Dhanasekaran, and S. Cobey. Lineage-specific protection and immune imprinting shape the age distributions of influenza B cases. Nature Communications, 12(1):4313, 2021. doi: 10.1038/s41467-021-24566-y.

